# An expanded reference catalog of translated open reading frames for biomedical research

**DOI:** 10.1101/2025.07.03.662928

**Authors:** Sonia Chothani, Jorge Ruiz-Orera, Jack A. S. Tierney, Jim Clauwaert, Eric W Deutsch, M. Mar Alba, Julie L. Aspden, Pavel V. Baranov, Ariel Alejandro Bazzini, Elspeth A. Bruford, Marie A. Brunet, Tristan Cardon, Anne-Ruxandra Carvunis, Claudio Casola, Jyoti Sharma Choudhary, Kellie Dean, Pouya Faridi, Ivo Fierro-Monti, Isabelle Fournier, Adam Frankish, Mark Gerstein, Norbert Hubner, Yunzhe Jiang, Manolis Kellis, Leron W. Kok, Thomas F Martinez, Gerben Menschaert, Pengyu Ni, Sandra Orchard, Xavier Roucou, Joel Rozowsky, Michel Salzet, Mauro Siragusa, Sarah Slavoff, Michal I Swirski, Nicola Ternette, Eivind Valen, Juan Antonio Vizcaino, Aaron Wacholder, Wei Wu, Zhi Xie, Yucheng T. Yang, Robert L. Moritz, Jonathan Mudge, Sebastiaan van Heesch, John R Prensner, Owen JL Rackham

**Affiliations:** Centre for Computational Biology, Cardiovascular and Metabolic Disorders, Duke-NUS medical school, Singapore 169857; Genome Institute of Singapore (GIS), Agency for Science, Technology and Research (A*STAR), 60 Biopolis Street, Genome, Singapore 138672, Republic of Singapore; Cardiovascular and Metabolic Sciences, Max Delbrück Center for Molecular Medicine in the Helmholtz Association (MDC), 13125 Berlin, Germany; European Molecular Biology Laboratory, European Bioinformatics Institute (EMBL-EBI), Wellcome Genome Campus, Hinxton, Cambridge CB10 1SD, UK; Department of Pediatrics, Division of Pediatric Hematology/Oncology, University of Michigan Medical School, Ann Arbor, MI, 48109, USA; Department of Biological Chemistry, University of Michigan Medical School, Ann Arbor, MI, 48109, USA; Institute for Systems Biology, Seattle, Washington, 98109, United States; Hospital del Mar Research Institute (HMRIB), Barcelona, Spain; 4Catalan Institute for Research and Advanced Studies (ICREA), Barcelona, Spain; School of Molecular and Cellular Biology, Faculty of Biological Sciences, University of Leeds, Leeds, LS2 9JT, UK; Leeds Omics, University of Leeds, Leeds, LS2 9JT, UK; Astbury Centre for Structural Molecular Biology, University of Leeds, Leeds LS2 9JT, UK; School of Biochemistry and Cell Biology, University College Cork, Cork, T12 K8AF, Ireland; Stowers Institute for Medical Research, 1000 E 50th Street, Kansas City, MO 64110, USA; Department of Molecular and Integrative Physiology, University of Kansas School of Medicine, Kansas City, KS 66160, USA; HUGO Gene Nomenclature Committee, Department of Haematology, University of Cambridge Clinical School, Cambridge CB2 0PT, UK; Medical Genetics Service, Pediatrics Department, University of Sherbrooke, J1E 4K8, Sherbrooke, Canada; Cancer Research Institute University of Sherbrooke, IRCUS, J1E 4K8, Sherbrooke, Canada; Centre de Recherche du Centre hospitalier universitaire de Sherbrooke, CRCHUS, J1E 4K8, Sherbrooke, Canada; Univ. Lille, Inserm, CHU Lille, U1192 - Protéomique Réponse Inflammatoire Spectrométrie de Masse - PRISM, F-59000 Lille, France; Department of Computational and Systems Biology, University of Pittsburgh School of Medicine, Pittsburgh PA 15260; Pittsburgh Center for Evolutionary Biology and Medicine (CEBaM), Pittsburgh PA 15260; Department of Ecology and Conservation Biology, Texas A&M University, College Station, TX, 77843 USA; The Institute of Cancer Research; School of Biochemistry and Cell Biology, University College Cork, Cork, Ireland T12 XF62; Centre for Cancer Research, Hudson Institute of Medical Research, Clayton, VIC, Australia; Monash Proteomics & Metabolomics Platform, Department of Medicine, School of Clinical Sciences, Monash University, Clayton, VIC, Australia; EMBL-EBI, Wellcome Genome Campus, Cambridgeshire, UK; Biozentrum, University of Basel, Switzerland; Institut Universitaire de France, ministère de l’Enseignement supérieur, de la Recherche et de l’Innovation, 1 rue Descartes, 75231 PARIS CEDEX 05, France; Program in Computational Biology and Bioinformatics, Yale University, New Haven, CT 06520, USA; Department of Molecular Biophysics and Biochemistry, Yale University, New Haven, CT 06520, USA; Department of Computer Science, Yale University, New Haven, CT 06520, USA; Department of Statistics and Data Science, Yale University, New Haven, CT 06520, USA; Department of Biomedical Informatics and Data Science, Yale University, New Haven, CT 06520, USA; Charité-Universitätsmedizin Berlin, 10117 Berlin, Germany; German Center for Cardiovascular Research (DZHK), partner site Berlin, Germany; Helmholtz Institute for Translational AngioCardiosciences (HI-TAC), Max Delbrück Center for Molecular Medicine at Heidelberg University, Heidelberg, Germany; MIT Computer Science and Artificial Intelligence Laboratory, Cambridge, MA, 02139, USA; Broad Institute of MIT and Harvard, Cambridge, MA, 02139, USA; Princess Máxima Center for Pediatric Oncology, Utrecht, 3584 CS, The Netherlands; Oncode Institute, Utrecht, The Netherlands; Department of Pharmaceutical Sciences, University of California, Irvine, Irvine, CA 92617, USA; Department of Biological Chemistry, University of California, Irvine, Irvine, CA 92617, USA; Chao Family Comprehensive Cancer Center, University of California, Irvine, Irvine, CA 92617, USA; BioBix, Lab for Bioinformatics and Computational Genomics, Faculty of Bioscience Engineering, Ghent University, Belgium; European Molecular Biology Laboratory, European Bioinformatics Institute (EMBL-EBI), Wellcome Genome Campus, Hinxton CB10 1SD, UK; Department of Biochemistry and Functional Genomics, Université de Sherbrooke, Sherbrooke, Canada; Institut Universitaire de France (IUF), 75000, Paris, France; Goethe University, Institute for Vascular Signalling, Centre for Molecular Medicine, Frankfurt am Main, Germany; CardioPulmonary Institute, Frankfurt am Main, Germany; Yale University Department of Chemistry, New Haven, CT 06520; Institute of Genetics and Biotechnology, University of Warsaw, Poland; University of Dundee, Dundee, UK; Department of Biosciences, University of Oslo, Oslo, Norway; European Molecular Biology Laboratory, European Bioinformatics Institute (EMBL-EBI), Wellcome Genome Campus, Hinxton, Cambridge, CB10 1SD, United Kingdom; Department of Computational and Systems Biology, School of Medicine, University of Pittsburgh, Pittsburgh, PA, 15213, USA; Pittsburgh Center for Evolutionary Biology and Medicine, School of Medicine, University of Pittsburgh, Pittsburgh, PA, 15213, USA; Singapore Immunology Network (SIgN), Agency for Science, Technology and Research (A*STAR), Singapore, 138648, Singapore; Department of Pharmacy & Pharmaceutical Sciences, National University of Singapore, Singapore, 117543, Singapore; State Key Laboratory of Ophthalmology, Zhongshan Ophthalmic Center, Sun Yat-sen University, Guangzhou, China; Institute of Science and Technology for Brain-Inspired Intelligence, Fudan University, 220 Handan Road, Shanghai 200433, China; School of Biological Sciences, University of Southampton, Southampton, UK

**Keywords:** GENCODE, Ribo-seq, TransCODE, non-canonical open reading frame

## Abstract

Non-canonical (i.e., unannotated) open reading frames (ncORFs) have until recently been omitted from reference genome annotations, despite evidence of their translation, limiting their incorporation into biomedical research. To address this, in 2022, we initiated the TransCODE consortium and built the first community-driven consensus catalog of human ncORFs, which was openly distributed to the research community via Ensembl-GENCODE. While this catalog represented a starting point for reference ncORF annotation, major technical and scientific issues remained. In particular, this initial catalogue had no standardized framework to judge the evidence of translation for individual ncORFs. Here, we present an expanded and refined catalog of the human reference annotation of ncORFs. By incorporating more datasets and by lifting constraints on ORF length and start-codon, we define a comprehensive set of 28,359 ncORFs that is nearly four times the size of the previous catalog. Furthermore, to aid users who wish to work with ncORFs with the strongest and most reproducible signals of translation, we utilized a data-driven framework (i.e. translation signature scores) to assess the accumulated evidence for any individual ncORF. Using this approach, we derive a subset of 7,888 ncORFs with translation evidence on par with canonical protein-coding genes, which we refer to as the Primary set. This set can serve as a reliable reference for downstream analyses and validation, with a particular emphasis on high quality. Overall, this update reflects continual community-driven efforts to make ncORFs accessible and actionable to the broader research public and further iterations of the catalog will continue to expand and refine this resource.

## Introduction

Since the creation of the Ensembl-GENCODE (hereafter GENCODE) project 20 years ago^1,2^, gene annotations have been broadly divided based on whether a given gene is *protein-coding* or *non-coding*. Today, the canonical set of coding sequences (CDS) produced by GENCODE is a key resource for understanding the translated portion of the transcriptome. However, an increasing understanding of the prevalence of translation outside the set of canonical protein annotations has revealed the presence of additional functional elements. Here, perhaps no development in genome technologies has been more provocative than the adoption of ribosome profiling (Ribo-seq)^3^, which has led to the identification of thousands of translated short unannotated open reading frames (typically 100 or fewer codons)^4–12^. We refer to these as translated non-canonical ORFs (ncORFs), specifically meaning ORFs with translation evidence that fall outside GENCODE’s standard protein-coding annotations. Hereafter, we omit the term ‘translated’ for brevity. In some cases, ncORFs are discovered to encode *bona fide* ‘microproteins’ that can then be classified as canonical^13^, leading to newly annotated protein-coding genes such as *MTLN*^14^*, MRLN*^15^, and *MIEF1-uORF/L0R8F8*^16–19^. However, many ncORFs may not meet the evolutionary constraints expected of canonical proteins^13,20^, yet potentially encode species-specific functional proteins^11,21^ or alternatively, serve regulatory roles through their translation^22–25^. NcORF translation has been shown to occur broadly under physiological conditions but there is great interest in the identification of ncORF translation products that may be recurrently or uniquely associated with specific cellular or pathological states^26–32^. Their widespread yet poorly understood nature highlights the need for standardized annotation to enable broader investigation by the scientific community.

Ultimately, the annotation of translated ncORFs depends on the accumulation and analysis of Ribo-seq data, the development of a new, more sophisticated classification schema for translation, and the additional inclusion of orthogonal protein evidence. To this end, in 2020, we initiated the global TransCODE Consortium, in collaboration with GENCODE^33^, HGNC (HUGO Gene Nomenclature Committee)^34^, UniProtKB (UniProt Knowledge-Base)^35^, PeptideAtlas^36^, and other leading academic labs. This consortium combines expertise in annotation, Ribo-seq, gene evolution, non-coding RNA function, and mass-spectrometry proteomics. The result of Phase I of TransCODE was an initial GENCODE-supported catalog of 7,264 ncORFs^20^ previously identified using Ribo-seq as well as guidelines on how to interpret Ribo-seq and other experimental data nominating such ncORFs for annotation^22^.

However, community-wide usage of these 7,264 ncORFs has revealed both their utility as well as their limitations. It has become clear that this first catalog of ncORFs contains blind spots, making an updated catalog essential for the community. Several such blind spots resulted from the criteria used for ncORF inclusion, which were employed at that time for practical, rather than biological, reasons. For example, the 7,264 ncORFs only include candidates that are: (1) greater than 15 codons in length, (2) initiated at AUG start codons, and, (3) the longest isoform in situations where multiple ncORFs share a large part of their amino acid sequence (equal to or greater than 90%). Yet, such filters likely hinder research on some ncORFs, particularly small ones. Indeed, it is known that ncORFs as small as two codons (i.e. ‘Start-Stop’ or ‘minimal’ ncORFs)^37^ can function as regulatory elements, and recent studies in human tissues and cell lines have demonstrated widespread translation of ncORFs below our previous size cutoff^11,12^. Moreover, translation products of ncORFs as short as eight codons can be presented on major histocompatibility complex molecules^27^. Similarly, non-AUG translation initiation events are increasingly recognized as biologically relevant, even within well-annotated protein-coding genes^38–40^, with increasing evidence that they are especially common within 5’ untranslated regions (5’ UTRs)^12,20^. Additionally, alternative splicing, the use of alternative initiation codons or sequence variation may be specific to disease states^21,26^ or even to different stages of the cell cycle^41^, making it possible to have multiple ncORF isoforms. Altogether, for the field of ncORF research to continue to grow, it is now critical to develop reference annotation of ncORFs further.

Here, we present an updated reference catalog of translated ncORFs detected by Ribo-seq based on GENCODE v45. As with the Phase 1 workflow, our goal was to use Ribo-seq data to map ncORFs to annotated transcripts, however, in this updated catalog we incorporate additional studies made available since the production of our first catalog and remove the restrictive filters (as described above, **Supplementary Fig. 1a, b**). We first generate a ‘Comprehensive set’ that substantially increases the number of ncORFs called from Ribo-seq from 7,264^20^ to 28,359. The new catalog adds thousands of ncORFs that were initially excluded from the first catalog, as well as ncORFs from the human body map translation project, extending the coverage of our resource to 11 additional primary human tissues and cell types^12^. We emphasize that *‘Comprehensive’* refers to the inclusive approach taken in incorporating ncORFs into this set, as opposed to the sense that the catalog may be biologically complete (which remains hard to assess). As another advancement over our previous work, we specifically designate ncORFs with robust evidence of translation in the pooled human body map Ribo-seq dataset into a ‘Primary set’ of ncORFs. To do so, we utilize a standardized assessment of ncORFs using translation signature scores that act as quality-control metrics, namely, ‘P-sites in Frame (PIF)’, ‘ORF Uniformity’, and ‘Dropoff’ scores (previously used in the human body map translation project^12^). Based on these scores, we denote a subset of 7,888 out of 28,359 ncORFs as the ‘Primary’ set, as these ncORFs have translation signature scores in the same range as canonical protein-coding sequences. The purpose of this Primary set (as opposed to the Comprehensive set) is to allow users to focus their analyses to ncORFs with the highest degree of translation evidence currently available. To ensure that these ncORF annotations can be widely applied across biomedical research, both sets are available at https://www.gencodegenes.org/pages/riboseq_orfs/.

## Results

### An expansion of the ncORF catalog

The new catalog of ncORFs is mapped to GENCODE v45 and is now referred to as the ‘v45’ catalog. It builds upon our initial release in July 2022, which was mapped to GENCODE v35^20^ (see **Table 1**). The v45 catalog incorporates datasets from the same seven published studies as used for the first catalog, and according to the same selection principles (**see Methods for details**) includes two additional human Ribo-seq ncORF datasets that were published in 2022^12^ and 2023^11^ (**Supplementary Table 1**). Firstly, Chothani *et al*^12^, carried out an extensive ncORF human translation body map study, generating more than a billion high-quality inferred P-site reads using 167 Ribo-seq samples. These samples covered six primary human cell types (atrial fibroblasts, coronary artery endothelial cells, umbilical vein endothelial cells, hepatocytes, vascular smooth muscle cells, embryonic stem cells) and five human tissues (brain, adipose tissue, heart, skeletal muscle, kidney), resulting in the identification of 7,767 ncORFs. Secondly, we incorporated 221 experimentally interrogated ncORFs reported by Sandmann *et al.*^11^, previously not included, as each was less than 16 codons in length. The v45 catalog also takes a more inclusive approach to ncORF identification; we now include ORFs smaller than 16 codons, those initiated by non-AUG codons, and all identified isoforms of each ncORF (**Supplementary Fig. 1a, 1b**).

**Table 1:** Comparison of v35 and v45 ncORF catalog.

|  | v35 catalog (2022) | v45 catalog (2025) |  |
| --- | --- | --- | --- |
|  |  | Comprehensive set | Primary set |
| Total ncORFs | 7,264 | 28,359 | 7,888 |
| Number of studies included | 7 | 9 | 9 |
| Number of Ribo-seq samples included | 139 | 226 | 226 |
| Overlap of v35 ncORFs | - | 7,091 | 1,024 |
| Length restriction | Yes (> 16 amino acids) | No | No |
| Start-codon restriction | Yes (AUG only) | No | No |
| Isoforms included | Partially (clustered isoforms ≥ 90% shared codon sequence) | Yes | Yes |
| Standardized metrics applied (Translation signature scores) using pooled body map data | No | No | Yes |

In total, we identified 28,359 ncORFs that we refer to as the ‘Comprehensive set’ of our v45 catalog (**Fig. 1a, Supplementary Table 2**). We retained most of the ncORFs (7,091 out of 7,264) from the first catalog, excluding 173 prior ncORFs because (i) they no longer mapped to GENCODE transcripts in v45 (n = 66), (ii) are now annotated as known proteins^13,20^ (n = 25), (iii) have been reassessed as extensions of known proteins (n = 80), or (iv) map to pseudogenes (n = 2) (**Supplementary Fig. 1c, Supplementary Table 3**). Building upon this, the Comprehensive set also includes: (1) 3,124 additional isoforms (due to alternative splicing or alternative translation initiation) of ncORFs that were present in the first catalog; (2) and a further 18,144 ncORFs that do not overlap with any of the ncORFs from the first catalog (**Supplementary Fig. 1d**).

**Figure 1.**
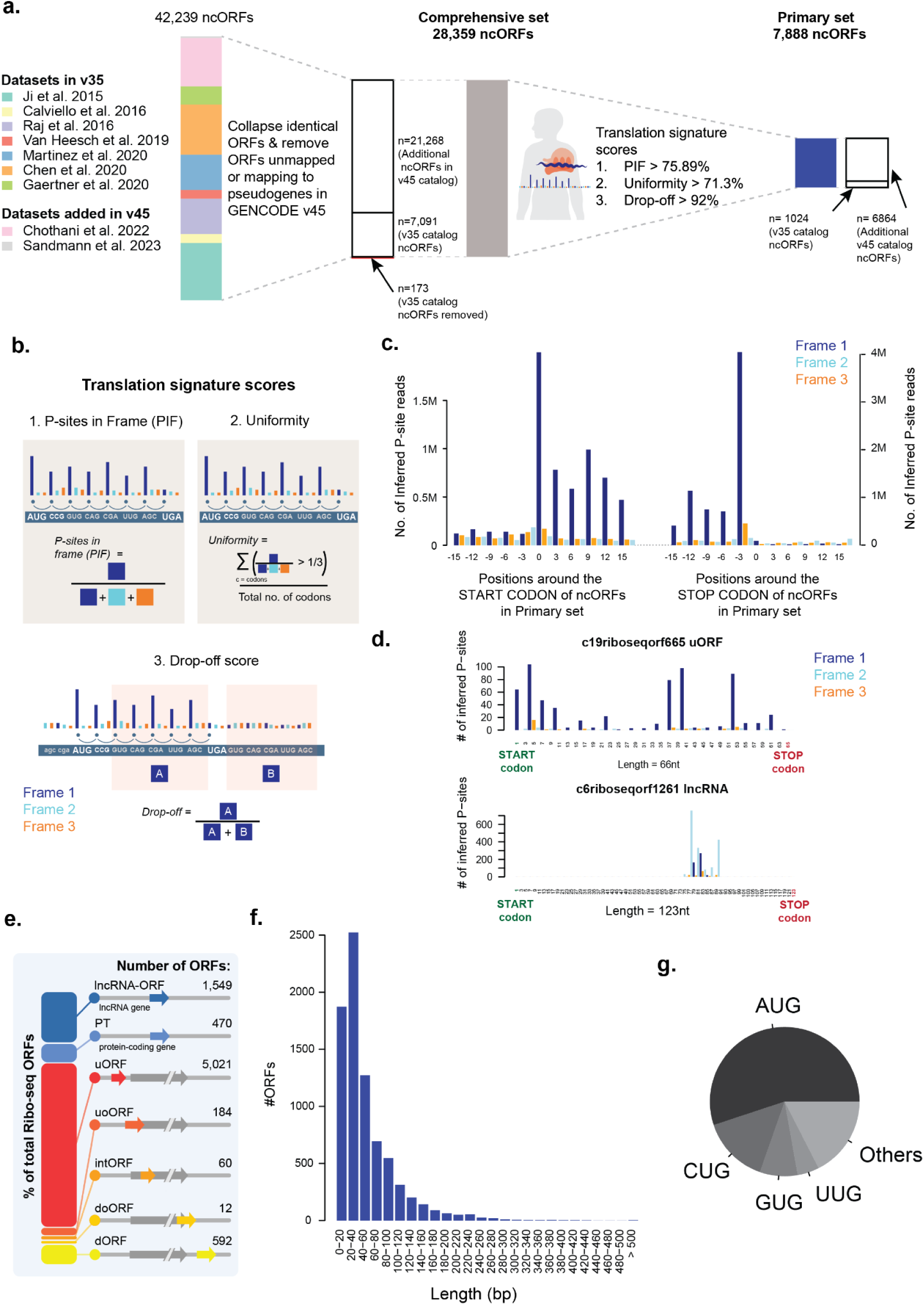
A Phase 2 GENCODE catalog for ncORFs using translation signatures. **a)** Sankey plot based flow chart showing number of ncORFs collected from nine studies^4–12^ without any hard filters on length or start-codon, followed by obtaining a comprehensive set by collapsing identical ncORFs and removing ncORFs not mapping to GENCODE v45 or mapping to pseudogenes and lastly the number of ncORFs in the *Primary* set obtained using translation signature scores. **b)** Translation signature scores and its quantification. This included three metrics to define translation signature presence in each ncORF. *P-sites in Frame (PIF)* quantifies the proportion of Inferred P-sites in the translating frame with respect to the total inferred P-sites in the ncORF. *Uniformity* quantifies the percentage of codons in the ncORF that have >33% inferred P-sites in the translating frame. *Drop-off score* quantifies the proportion of inferred P-sites before the stop-codon with respect to the total inferred P-sites in the translating frame in a 30bp window around the stop codon. **c)** A bar plot showing an aggregated P-site profile around the start- and stop-codons of the *Primary* set of ncORFs **d)** A bar plot showing P-site profile of two selected Ribo-seq ncORFs with high and low translation signature scores. **e)** A stacked bar chart showing numbers of *Primary* set ncORFs across different ncORF types. **f)** A histogram showing the length distribution of *Primary* set ncORFs. **g)** A pie chart showing the distribution of start-codons identified for the *Primary* set ncORFs.

### Data-driven discrimination of a Primary set of ncORFs for improved end-user usability

In gene annotation, filtered subsets help users navigate large transcript collections, such as the 19,252 Matched Annotation from NCBI and EBI (MANE) transcripts within the 252,989 total transcripts in GENCODE v45^42^. While MANE prioritizes functionally relevant, well-supported models, ncORFs currently lack equivalent functional information. Therefore, we rely on data-driven metrics to identify a filtered set of ncORFs. In our initial catalog (v35)^20^, we tested for replication of ncORFs across multiple studies as a proxy for detection reliability (**Supplementary Fig. 1e-g**). However, this approach is constrained by substantial variability in both the quality of the individual Ribo-seq dataset as well as the nature and performance of computational workflows deployed. These issues are further confounded by the relative sparseness of Ribo-seq reads across an ncORF when using single samples or smaller Ribo-seq datasets^13,20,43^. To address these inconsistencies, here we define a Primary set by using a large pooled human body map dataset to quantify numerous data-driven metrics that each ncORF must satisfy, irrespective of the method of identification, thereby ensuring a more robust and consistent filtering strategy.

To construct the Primary set, we apply filters based on features derived from Ribo-seq data, selecting ncORFs whose translation signatures closely resemble those of canonical protein-coding genes. As a result, to standardize the evidence for ncORFs obtained from different studies, we independently assessed each ncORF to identify a subset of ORFs featuring the same degree of experimental evidence as observed for canonical CDSs. Specifically, we used Ribo-seq data-derived P-site profiles using the pooled human body-map data to assess each individual ncORF in the Comprehensive set using previously designed metrics^12^: i) ‘P-sites in Frame (PIF)’, which determines the proportion of inferred P-site reads that are found in the translating frame; ii) ‘Uniformity’, which quantifies the proportion of codons within the ncORF that have more than 33% reads in the translating frame. iii) the ‘Drop-off score’, which quantifies the proportion of P-sites in-frame before the stop codon to the total P-sites in frame within the 30 bp window before and after the stop codon. An ncORF from the Comprehensive set was included in the Primary set if it passed a threshold of 75.9% for PIF, 71.3% for Uniformity, and 92.0% for Drop-off. These thresholds were previously determined from annotated CDSs^12^ using the pooled human bodymap data to identify ncORFs with similar translational signatures (**Fig. 1b, see Methods for details**). As a result, we compiled a Primary set of 7,888 ncORFs (from the 28,359 in the Comprehensive set) (**Fig. 1a, Supplementary Table 4,** comparison to v35 release shown in Fig. 1a and Table 1). On aggregating P-sites around the start and stop of the ncORFs, we found clearer translation signatures for the Primary set as compared to the remainder of the Comprehensive set (**Fig. 1c-d, Supplementary Fig. 1h, 2**).

### ncORFs in the Primary set with alternative features

Of the 7,888 ncORFs in the Primary set, 5,027 ncORFs (63.7%) are shorter than 16 codons (**Fig. 1f, Supplementary Fig. 1k**), and 3,543 ncORFs (44.9%) start from non-AUG initiation codons (**Fig. 1g**), two thresholds applied in our initial v35 catalog. In the Primary set, we found that almost two-thirds of the identified ncORFs are upstream ORFs (uORFs, n = 5,021, 63.7%, **Fig. 1e**) followed by ORFs in long non-coding RNA (lncRNA) genes (lncRNA-ORFs, n = 1,549, 19.6%) (Potential explanations of these observations are discussed below in the Limitations section). For example, the Primary set includes the 3-codon-long uORF c22riboseqorf449 in *ATF4*, which is known to influence downstream CDS translation upon stress or disease^44^. The Primary set also includes 69 ‘minimal ORFs’ that only contain a start and a stop codon. Such minimal ORFs could serve as regulatory elements^37^ and thus an important inclusion for reference annotations^45^. Within the Primary set, we found that 55.1% of ncORFs initiate at AUG codons, 14.5% at CUG and 8.2% at GUG, similar to what has been previously reported by others^46^ (**Fig. 1g**).

Next, we sought to define ncORF isoforms in the Primary set that can arise via alternative splicing and/or alternative initiation. These processes have historically been challenging to resolve in Ribo-seq data due to the sparsity of Ribo-seq coverage throughout the length of individual ncORFs^22^, as well as complexities in the underlying transcript models. Furthermore, it is also expected that both processes can be subjected to differential expression, i.e., across cell types and different biological conditions. Within the Primary set, we identified 2,247 ncORF isoforms that partially share the amino acid sequences of other ncORFs within the same set (**Supplementary Fig. 3a**), representing 28.4% of the total set. Among these, 438 ncORFs shared the same start position, whereas 428 shared the same stop position with another ncORF (**Supplementary Fig. 3a**). Notably, only 258 ncORFs exhibited an overlap of ≥ 90% with other ncORFs, indicating that most ncORF isoforms contributed substantial stretches of unique codon sequences. We also identified 868 non-AUG initiating ncORFs as isoforms of AUG ncORFs and 2,675 non-AUG initiating ncORFs that do not have other AUG-initiating ncORFs included in the Primary set. For instance, *LMBRD2* encodes two overlapping uORFs with partially shared codon sequences that are translated into two small ncORF products of 38 and 41 codons, respectively, in the complete absence of a nearby AUG (**Supplementary Fig. 3b**). As another example, the *GIHCG* lncRNA contains four different ncORF variants starting with a mix of alternative non-AUG and AUG codons that each display convincing evidence of translation initiation (**Supplementary Fig. 3c**).

### Limitations: Assessment of gaps in the Primary set

Not all ncORFs included in our catalog were identified using the tissues or primary cells used in the human body map Ribo-seq dataset. However, we used this extensive dataset to score ncORFs based on translation signature scores for inclusion into the Primary set. To address this potential blind spot in our workflow, we tested whether the human body map data provided sufficient coverage to test all ncORFs, regardless of their source tissue or cell type. Evaluating each ncORF for their expression levels, we found that 78.2% of ncORFs in the Comprehensive set (22,179 out of 28,359) had a P-site Per Million (PPM) > 1, allowing for their confident analysis. In contrast, 6,180 ncORFs had low P-site coverage (PPM < 1); despite their low expression ∼16% still passed the translation signature scores and met the criteria for inclusion in the Primary set (**Fig. 2a**). However, it is important to note that some ncORFs with low coverage that did not meet the criteria may exhibit high translation scores in other tissues or cell lines that were not analyzed in this study.

**Figure 2.**
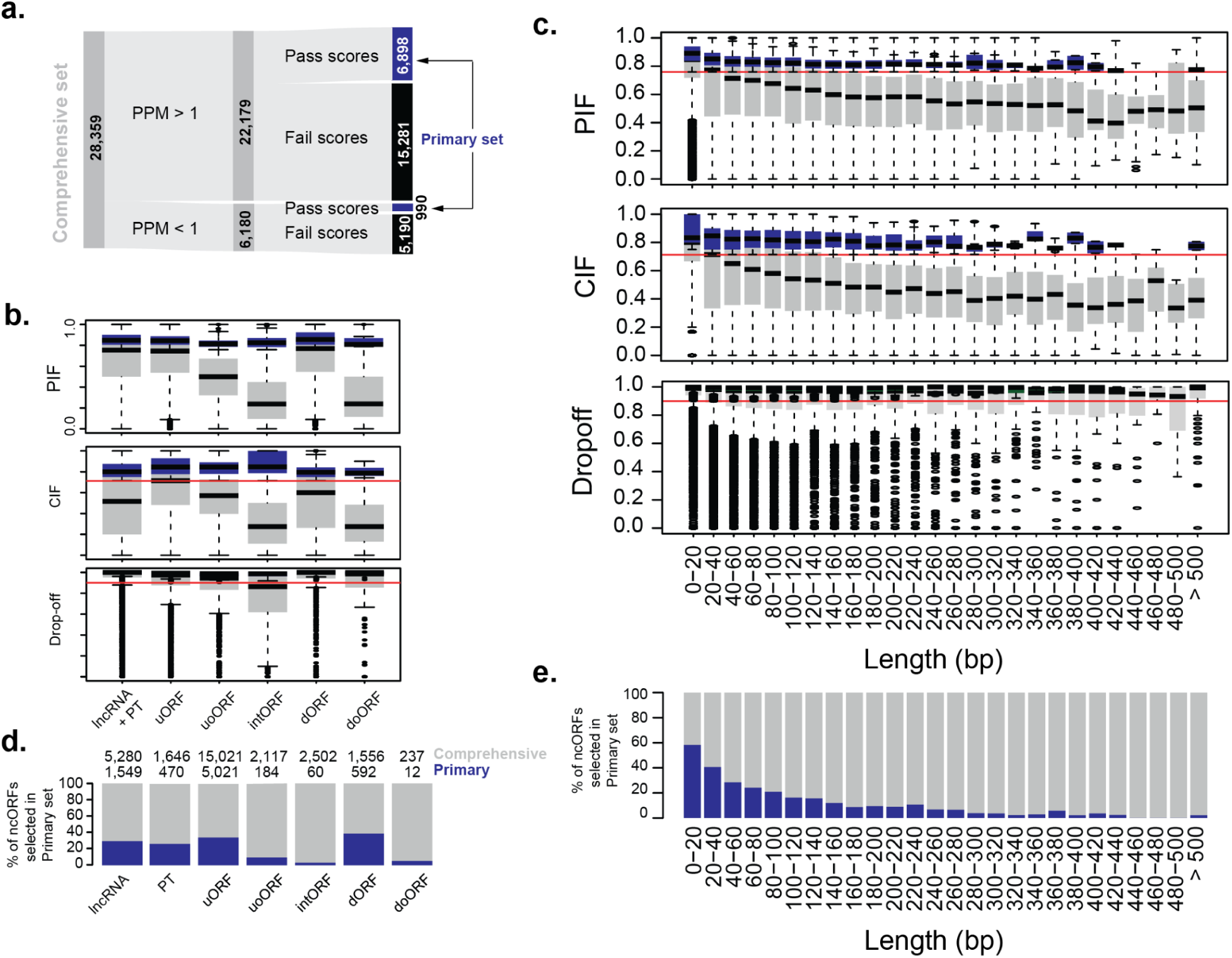
Developing translation signature score as confidence levels for ncORFs. **a)** A Sankey plot showing the number of ncORFs in the comprehensive set and corresponding ORFs that have P-sites Per Million or PPM > 1 in the pooled human body-map Ribo-seq data followed by the number of ncORFs that pass the translation signature scores. **b)** Boxplot showing the distribution of translation signature scores (P-sites in Frame (PIF), Uniformity, and Drop-off scores) across various ncORF types. **c)** Boxplot showing the distribution of translation signature scores across various length bins of ncORF length in base pairs. **d)** Stacked barplot showing the proportion of ncORFs passing thresholds within each ncORF type (**d**) and length bin (**e**). Gray: Comprehensive set ncORFs, Blue: Primary set.

Furthermore, we tested whether ncORF length and type (**Fig. 1e**, **see Methods for details**) influenced their chances of being included in the Primary set using the abovementioned Ribo-seq quality metrics. We did so because we reasoned that metrics like Uniformity could be biased in favor of short ncORFs, where ribosome footprint coverage across a small number of codons would still result in good uniformity. Similarly, we anticipated that ncORF types that overlap other ncORFs in alternative reading frames would score poorly on each of the three metrics. Indeed, we found that smaller ncORFs perform relatively well compared to longer ncORFs, indicating a length-dependent reduction in ncORF performance for the PIF and Uniformity metrics (**Fig. 2b**). For example, 48.4% (9,384 out of 19,382) of ncORFs <100 nucleotides in length had a PIF score above the threshold, compared to 24.6% (2,205 out of 8,977) of ncORFs >100 nucleotides (Chi-square test, p-value 1.95 x 10^-143^). Secondly, we found that ncORFs overlapping with CDS sequences in a different reading frame, such as upstream overlapping ORFs (uoORFs), downstream overlapping ORFs (doORFs), or internal ORFs (intORFs) had limited representation in the Primary set as the scores assume a single ORF is translated in the given region (**Fig. 2c, Supplementary Fig. 3d, e**). Together, this shows that longer ncORFs, overlapping ORFs, and internal ORFs may be under-represented in the Primary set, and investigators interested in these ncORF types can source these from the Comprehensive set instead.

### Availability of the new GENCODE catalog

The Comprehensive and Primary sets are available at https://www.gencodegenes.org/pages/riboseq_orfs/. Regardless of their current level of supporting data, we emphasize that every ncORF in the Comprehensive set has been published as being translated in at least one of the nine studies included, and thus serves as a valuable reference for tracking and comparison in ncORF discovery, even if absent from the Primary set. We anticipate that some Comprehensive set ncORFs may gain additional supporting data over time, allowing them to be reclassified into the Primary set in future updates.

## Discussion

Despite significant advances in our understanding of the human genome, gene and ORF annotations remain a work in progress. Redundancy in independent efforts for generating ncORF catalogs, as well as low overlap across published sets, has highlighted the need for a unified reference annotation. This need has been further amplified by ongoing efforts to characterize potential protein-coding genes from ncORF catalogs^13,47^. Here, we have assembled an updated catalog of human ncORFs, seeking to address several current gaps in ncORF annotation. We now provide an expanded Comprehensive set of 28,359 ncORFs, aggregated from the nine studies included in this work, without any length or start-codon filters (considering all possible start codons reported in their respective studies), intended for efforts aiming to assess the ncORF search space in an inclusive manner. Additionally, we denote a Primary set of 7,888 ncORFs that have clear translation signatures in the pooled human bodymap data using a combination of PIF, Uniformity, and Drop-off scores. This subset may be useful for analyses where limiting false positives is critical, and we expect these highly supported ncORFs will serve as a ‘gold standard’ set. We view the value of this updated catalog as threefold: (1) a unified resource for researchers; (2) a high-quality Primary set that may serve as a field reference; and (3) the development of standardized metrics to quantify translation evidence. These features may prove valuable for worldwide efforts to characterize the biological function of ncORFs^8,26,48–50^, the protein-coding potential of ncORFs^13^ including initiatives such as the Understudied Proteins Initiative^51^, iMOP^52^, the evolutionary dynamics of ncORFs^11,21,53^, the tissue- and disease-specificity of ncORFs, and the potential for nucleotide variants to impact ncORFs in human health and disease.

Potential users of this updated ncORF catalog should note that the Primary set has many very short ncORFs; 56.2% of all annotated ncORFs are less than 16 codons in length. In part, this is to be expected from both a technical and biological perspective. Shorter ncORFs can achieve high PIF and Uniformity scores with fewer base pairs tested, and detection of shorter ncORFs may also be expected, given that longer ncORFs are more likely to trigger nonsense-mediated decay^54^. Many of these short ncORFs are found in the 5′ UTRs of mRNAs, and because mRNAs have higher transcript expression overall compared to lncRNAs, this may increase detection sensitivity for uORFs and uoORFs^55^. Additionally, 5′ UTRs are known hotspots for the emergence of microproteins and play a central role in translational regulation, including canonical protein expression control^56,57^. An inverse relationship between uORF length and translational reinitiation efficiency has also been observed^23^ and recent population genetics studies provide further support for the functional relevance of these elements through evidence of sequence constraint^58^. Therefore, the enrichment of short uORFs in the Primary set is likely driven by biological factors – such as the overall higher level of translation for uORFs compared to lncRNA-ORFs, downstream ORFs (dORFs), and others^26^ – rather than false positives from the original ORF callers or artifacts from experimental protocols, as the included datasets are not biased to incorporate cells treated with antibiotics (e.g. homoharringtonine, lactimidomycin) that stall ribosomes in the 5’UTR^59,60^. Another key feature in this catalog is the inclusion of various isoforms of a given ncORF, such as different splicing isoforms or alternative start-or stop-positions. We included these for three main reasons. First, they likely reflect genuine transcriptional and translational complexity rather than technical artifacts. Second, comprehensive annotation supports efforts to characterize translation and enables tracking of isoforms that may be differentially detected across studies, because different RNA isoforms can expose alternatively translated ncORFs whose expression varies across cell types, tissue, or conditions. Third, given the current limitations in assigning function, excluding isoforms prematurely may overlook biologically relevant candidates not captured in the Primary set.

Nevertheless, we acknowledge the limitations of this work. First, our usage of published ncORF calls from a limited number of publications may have a bias towards ncORFs identified in specific samples, by specific research groups, or by certain Ribo-Seq analysis tools/protocols. Second, trusting the ncORF calls made by other publications necessitates accepting decisions made about pre-processing and post-processing steps of the data, which may differ between research efforts. Notably, recent machine learning approaches employ no pre-processing steps for the data^61^ or use sequence information to improve the prediction of Ribo-seq signal^62–64^. Third, scoring metrics such as PIF and Uniformity are biased towards certain ncORFs, particularly small ones (as noted above) and biased against those overlapping with other translated reading frames or CDSs. In overlapping ncORFs, periodic ribosome footprint signals become mixed, making their independent evaluation difficult and leading to an underestimation of their presence^22^. Fourth, start-codon determination and assigning the correct RNA isoform to each ncORF remains challenging. Lastly, here, we refer to these translation regions as ncORFs, noting that this is not a codified term in our databases and we anticipate a new ontology system will be developed in due course. For all of these reasons, we regard reference annotation to be an ongoing and iterative process and hope to be able to better address these potential concerns in future catalog iterations as more Ribo-seq datasets are incorporated and methods for ncORF detection improve.

Although the Primary set is unlikely to represent a complete survey of ncORFs with biological relevance, we expect the filtering approach taken here increases the rate of accurate ncORF detection and provides a subset of ncORFs with the highest utility for the life sciences and medical communities. Moreover, our resource provides a comprehensive overview of Ribo-seq evidence for all ncORFs included in both the Primary and Comprehensive sets. Researchers can customize their analyses by selecting their own thresholds for PIF, Uniformity, and Drop-off scores, or by filtering based on additional metadata - such as expression levels or ORF length. This flexibility enables optimization for various downstream applications, including mass spectrometry, immunopeptidomics^13^, evolutionary analyses^11,13^, testing for degradation signals^65,66^, interaction network mapping^67^, and other strategies aimed at identifying functional candidates aligned with specific research goals.

In summary, we present here an expanded catalog of ncORFs aligned with the GENCODE v45 resource and support a Primary set of 7,888 ncORFs that have high-quality evidence of their translation. We believe that this work will be widely applicable to future biomedical research on ncORFs in diverse settings, including human health and disease.

## Supplementary Tables

Note: This list has not been subject to peer review Filename: Supplementary_Tables.xlsx

## Acknowledgments

S.C was supported by Khoo post-doctoral fellowship award, SingHealth and Singapore Ministry of Health’s National Medical Research Council under Open Fund - Young Individual Research Grant (OFYIRG23jan-0034/MOH-001340). E.W.D and R.L.M were supported in part by the National Institutes of Health grants R24 GM148372 (E.W.D.), R01 GM087221 (E.W.D., R.L.M.), S10 OD026936 (R.L.M.), and by National Science Foundation grants DBI-2324882 (E.W.D.) DBI-1933311 (E.W.D.), and MRI-1920268 (R.L.M.). N.H was supported by an ERC-advanced grant under the European Union Horizon 2020 Research and Innovation Program (AdG788970), the Deutsche Forschungsgemeinschaft (German Research Foundation - DFG CRC/SFB-1470 – B03), the Pathfinder Cardiogenomics Programme of the European Innovation Council of the European Union DCM-NEXT, and in part by a grant from the Chan Zuckerberg Foundation. E.B. acknowledges funding from the US National Institutes of Health (NIH) National Human Genome Research Institute (NHGRI) under award number U24HG003345. J.A.V. is supported by funding from Wellcome [grant number 223745/Z/21/Z], and from EMBL core funding. J.M.M. and A.F are supported by the Wellcome Trust (grant number 108749/Z/15/Z), the National Human Genome Research Institute (NHGRI) of the US National Institutes of Health (NIH) under award number 5U24HG007234 and the European Molecular Biology Laboratory (EMBL). The content is solely the responsibility of the authors and does not necessarily represent the official views of the National Institutes of Health. Ensembl is a registered trademark of EMBL. S.v.H. acknowledges funding from Fonds Cancers (FOCA, Belgium), Stichting Reggeborgh (the Netherlands), and Villa Joep. This publication is part of the project “Evolutionarily young microproteins in childhood brain cancer” (with project number VI.Vidi.223.022 of the research programme NWO talent programme Vidi, which is (partly) financed by the Dutch Research Council (NWO), awarded to S.v.H. Research reported in this publication was supported by Oncode Accelerator, a Dutch National Growth Fund project under grant number NGFOP2201, awarded to S.v.H. This work is co-financed by Oncode Institute, which is partly funded by the Dutch Cancer Society. J.R.P. acknowledges funding from the National Institutes of Health / National Cancer Institute [K08-CA263552-01A1]; the V Foundation for Cancer Research [V2024-013]; Tough2gether Foundation; Hyundai Hope on Wheels Foundation; the Yuvaan Tiwari Foundation; DIPG/DMG Research Funding Alliance; Book for Hope Foundation; CureSearch for Cancer Research; Morgan Adams Foundation; the Lindonlight Collective [GR-24-008 and GR-24-012] the Chad Carr Pediatric Brain Tumor Center; and the University of Michigan Rogel Cancer Center. J.R.P. is the Ben and Catherine Ivy Foundation Clinical Investigator of the Damon Runyon Cancer Research Foundation [CI-127-24].

## Methods

### Collecting ncORFs from multiple datasets and mapping to GENCODE v45

We previously initiated the first catalog by selecting seven distinct Ribo-seq ncORF datasets from a range of human studies, each of which has been pivotal for genome-wide ncORF identification using Ribo-seq over the past decade^20^. Here, we included these seven datasets plus two new Ribo-seq ncORF datasets that were published in 2022^12^ and 2023^6^ (**Supplementary Table 1**).

When available, we retrieved exonic coordinates and sequences for these ncORFs. For datasets based on the older Human Genome assembly (GRCh37/hg19), we converted ORF coordinates to GRCh38/hg38 using UCSC Liftover. We compiled a total of 42,239 ncORFs. All translated ORF sequences were remapped to the Ensembl Release v.111 transcriptome (equivalent to GENCODE v45), collapsing identical genomic ncORF sequences. This resulted in 30,103 unique genomic sequences after excluding 1,452 ORFs that could not be matched to any transcript and 10,684 identical ncORF regions. Our selection was limited to ncORFs found in lncRNAs, non-coding RNA transcripts within known protein-coding genes, alternative reading frames of CDS annotated within known protein-coding genes, and untranslated regions (UTRs) of coding transcripts annotated within known protein-coding genes. Therefore, we excluded 1,744 ncORFs partially or totally overlapping annotated CDSs in the same frame (protein-coding or nonsense mediated decay) or pseudogenes in any frame, as was done for the previous catalog^20^. These exclusions were necessary as the ncORF datasets were based on older transcriptome versions, and newly annotated protein-coding sequences and pseudogenes are now available in GENCODE v45. Of 80 v35 catalog ncORFs that are now reclassified as CDSs, 49 previously overlapped incomplete protein-coding CDSs annotated with missing 5’ or 3’ regions but we now completely classify these cases as CDS. Pseudogenes were removed because not all the original studies included only uniquely mapped reads, which could have led to false identifications of such sequences, although we acknowledge that many pseudogenes are known to be expressed and translated.

In total, we generated a Comprehensive set of 28,359 ncORFs. Of these, 7,091 ncORFs were already part of the first catalog and 21,268 were newly added. Unlike the first catalog, we did not apply any minimum length filter or exclude non-AUG start codons. Non-AUG ncORFs were predicted in five of the nine considered studies, with the exception of those using RiboTaper or ORFquant, which only annotated AUG-initiated ncORFs (van Heesch et al. 2019, Calviello et al. 2016, Gaertner et al. 2020, Sandmann et al. 2023). Notably, in Martinez et al. 2020, ncORFs without an identifiable AUG start codon were defined from stop codon to stop codon, facilitating the inclusion of non-AUG-initiated ncORFs. Because of this, we acknowledge that some of the non-AUG ncORFs presented here may have inaccurate initiation site annotations.

### Transcript assignment and classification of identified ncORFs

The vast majority of ncORFs overlapped several transcript models within a given gene and could not be uniquely mapped to a single host transcript. To resolve these ambiguities, we assigned the ncORF to the main isoform selected using MANE^42^. For the rest of the cases that could not be assigned to the MANE Select isoform, we chose the isoform with the highest APPRIS^68^ score as the most probable isoform translating the ORF. In cases where multiple transcripts had comparable APPRIS scores, we further examined the Ensembl transcript support level (TSL) scores and selected the one with the highest support, prioritizing protein-coding transcripts over non-coding ones. All potential Ensembl transcript and gene IDs associated with each ORF, as well as the selected host transcripts, are detailed in **Supplementary Table 2**. While we made sure that the ncORFs do not partially or totally overlap the amino acid sequences of any annotated CDSs in the same region, it is possible that future releases of transcript annotations will include new models, and at least some ncORFs may be reannotated as new CDS extensions resulting from alternative splicing^69^.

NcORFs were categorized into seven distinct types exactly as defined for the first catalog^20^, i.e. based on the biotype of the assigned host isoform and the ORF position relative to known canonical protein-coding sequences.

**lncRNA-ORFs**: ncORFs found on long non-coding RNA genes.

**PT-ORFs**: ncORFs encoded by non-coding transcripts from protein-coding genes.

**Upstream ORFs** or **uORFs**: encoded within 5’ UTR sequences.

**Upstream overlapping ORFs** or **uoORFs**: encoded within 5’ UTR sequences and partially overlapping an already annotated CDS downstream in an alternative frame.

**Internal ORFs** or **intORFs**: completely overlapping within an already annotated CDS in an alternative frame.

**Downstream ORFs** or **dORFs**: encoded within 3’ UTR sequences

**Downstream overlapping ORFs** or **doORFs**: encoded within 3’ UTR sequences and partially overlapping an already annotated CDS in an alternative frame.

Lastly, as a further advancement in this catalog, we now modify our GENCODE naming of these ORFs. We have decided not to proceed with the dual system used previously, e.g. c1riboseqorf1 / c2norep2. This decision was made because we consider that ORFs should not be named according to their reproducibility, e.g. c2norep2, as this could potentially change. Therefore, for this catalog, we now use a simplified system based around only ‘c1riboseqorf1’, ‘c1riboseqorf2’. We keep the names of ncORFs from the previous catalog where appropriate, i.e. for those that were called as replicated, and subsequently continuing numbering ‘upwards’ to assign i) new names for ncORFs found in the previous catalog that were not replicated (i.e. previously annotated as ‘norep’) and ii) names for novel ncORFs not included in the first catalog. The ‘norep’ names that were used in the first catalog are also listed in the new datafile as ‘legacy_names_v35’ to allow comparison with the previous catalog. We anticipate that a standarised nomenclature system classifying for ncORFs will be devised in due course.

### Isoforms of ncORFs and overlap across studies

We compiled ncORFs from various studies, each using different datasets and methodologies. While identical ncORFs were treated as unique instances, we also identified many cases of ncORFs sharing part of their codon sequences. Therefore, ncORFs sharing any codon sequence were classified as ncORF isoforms. All isoforms were included in both the Comprehensive and Primary sets.

### Quantifying translation signature scores for ncORFs

We quantified translation signature scores for each ncORF in the Comprehensive set to test for evidence of translation in our previously pooled large-scale human body-map Ribo-seq data. The pooled dataset was obtained using the same workflow as previously described. Read alignment was carried out using STAR^70^ and only uniquely mapped reads were retained. From these, only reads with lengths between 27–30 nucleotides and exhibiting >60% overall 3-nt periodicity in annotated protein-coding genes were selected to generate the P-site read files. P-site files from individual samples were then combined to create the pooled body map dataset, which was used to quantify translation signature scores.

Translation signature scores include three metrics as previously described^12^, namely 1) P-sites in Frame (PIF), 2) Uniformity and 3) Drop-off. P-sites in Frame were calculated as the proportion of inferred P-site reads in the translating frame of the ncORF to the total inferred P-site reads in the ncORF. For the Uniformity calculation, each codon was tested for the proportion of P-site reads in Frame 1 to the total reads in the codon. The number of codons that had more than 33% of reads in Frame 1 were considered as having evidence for translation and the proportion of codons that show translation to the total number of codons was considered as the Uniformity. Lastly, for Drop-off score, a 15bp transcript flanking window on either side of the stop-codon was selected. P-site reads before and after were quantified and Drop-off was calculated as ratio of reads before the stop-codon with respect to the total reads in both before and after flanking regions. These three scores were calculated for each ncORF in the Comprehensive set. These scores were also quantified previously^12^ for known protein-coding ORFs to identify ncORFs with similar translation signatures. A normal distribution was fitted to the score distributions of annotated protein-coding ORFs and thresholds were defined based on the 95th percentile: 75.89% for PIF, 71.3% for Uniformity and 92% for Drop-off. The ncORFs in the Comprehensive set that passed these thresholds were considered to have high evidence for translation and selected for the Primary set. The ncORFs in the Primary set and the Comprehensive set were ranked for the Uniformity scores and top 10 ncORFs from Primary set and bottom 10 ncORFs from Comprehensive set were selected for visualization (P-sites per million or PPM < 1 were skipped in the selection. PPM calculation described in the next section). The scores were able to select for individual ncORFs which showed clear periodicity throughout the length of the ncORFs, thus discriminating ncORFs from ncORFs with just a single read stack or discontinuous periodicity or periodicity in the wrong frame (**Supp. Fig. 2**).

### Testing limitations in Primary set

P-sites per million (PPM) values were calculated by normalizing ORF P-site counts by the ORF length and scaling to per-million total normalized counts. PPM values were quantified for each ncORF and annotated protein-coding ORF and PPM > 1 values were considered as a threshold for expression (**Fig. 2a**). NcORFs in the Comprehensive set and Primary set were split into length bins of 20 nucleotides (nt) each until 500nt and into ncORF type categories such as uORF, dORF, uoORF, doORF, lncRNA-ORF, PT-ORF and scores for such length bins and categories were presented using boxplots. Proportion of ncORFs in each bin and category were quantified and a chi-square test was carried out to test if the proportion of ncORFs that passed the PIF and Uniformity scores changed for ncORFs that are less than or more than 100nt.

---

**Supplementary Figure 1.**
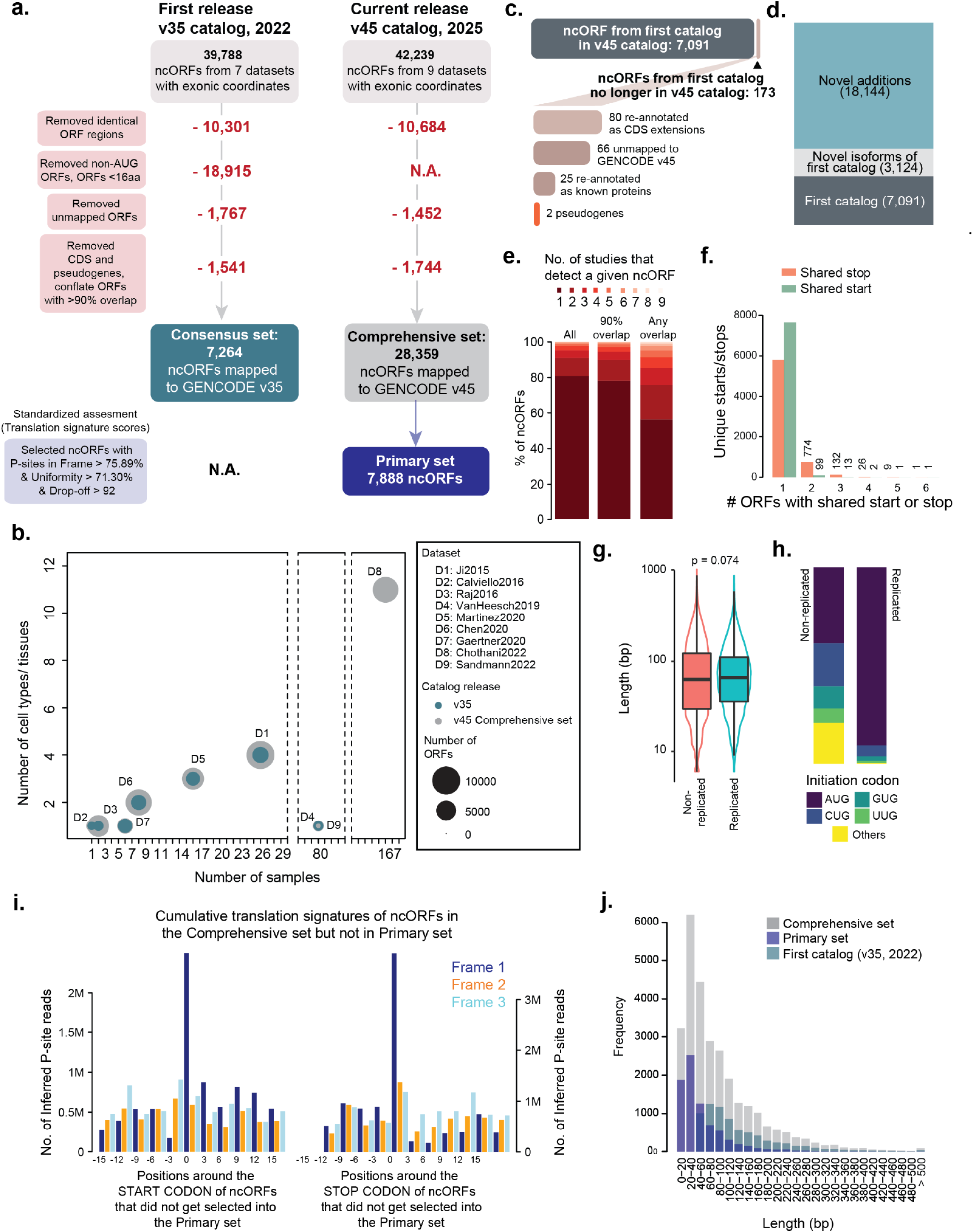
**a)** A schematic of the filtering steps for the previously published catalog in 2022 (v35) and the current v45 catalog. Numbers in red denote ncORFs that were removed. **b)** A bubble plot showing number of cell types or tissues vs number of samples sequenced with circle size denoting the number of ncORFs identified in a given study. Light blue: NcORFs retained in v35 catalog. Gray: NcORFs retained in v45. **c)** Number of ncORFs from the v35 catalog that were retained/not retained in the updated v45 catalog. Re-annotated to CDS includes cases that overlapped annotated proteins with missing 5’ or 3’ ends, or newly annotated extensions which have now been updated on GENCODE. **d)** Stacked barplot showing number of ncORFs included in the v45 catalog that were also included in the first catalog (v35 catalog, n=7,091), new additions that are isoforms of ncORFs included in the first catalog (n=3,124), and novel additions to this v45 catalog (n=18,144). **e)** Stacked bar plot illustrating the overlap of ncORF sequences identified across nine studies. The analysis includes all individual ncORFs (left), ncORFs clustered based on 90% sequence overlap (middle), and ncORFs grouped by any degree of overlap (right). **f)** Barplot showing a summary of isoforms within the v45 catalog. Orange: ncORFs have the same stop-coordinate, Green: ncORFs have the same start-coordinate. **g)** Box plot showing the length differences between replicated and non-replicated ncORFs. Wilcoxon test, p-value = 0.074. **h)** Stacked bar plot illustrating the proportion of initiation codons in replicated and non-replicated ncORFs. **i)** P-site profiles around the START and STOP codon of ncORFs in the Comprehensive set that did not get selected into the *Primary* set. Dark blue: P-sites in Frame 1, Orange: P-sites in Frame 2, Light blue: P-sites in Frame 3. **j)** Barplot showing length distribution of ncORFs. Gray: v45 catalog, Comprehensive set, Dark blue: v45 catalog, Primary set, Light blue: first catalog (v35 catalog, 2022).

**Supplementary Figure 2:**
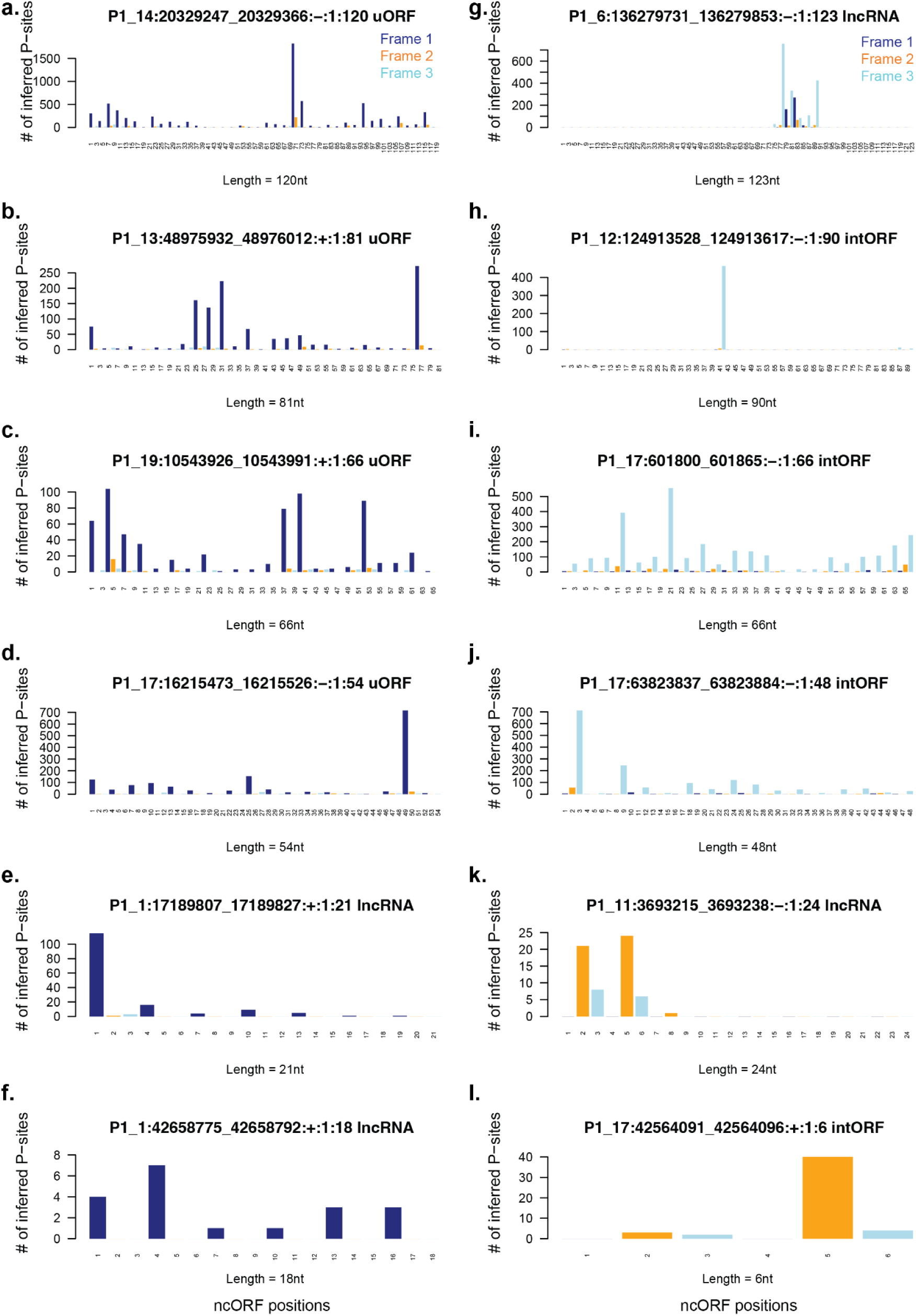
Individual examples of high-scoring and low-scoring ncORFs. **a-f.** Barplots showing number of P-sites throughout the length of high-scoring ORFs. **g-l.** Barplots showing number of P-sites throughout the length of low-scoring ORFs. Dark blue: Translating frame/Frame 1, Orange: Frame 2, Light blue: Frame 3.

**Supplementary Figure 3.**
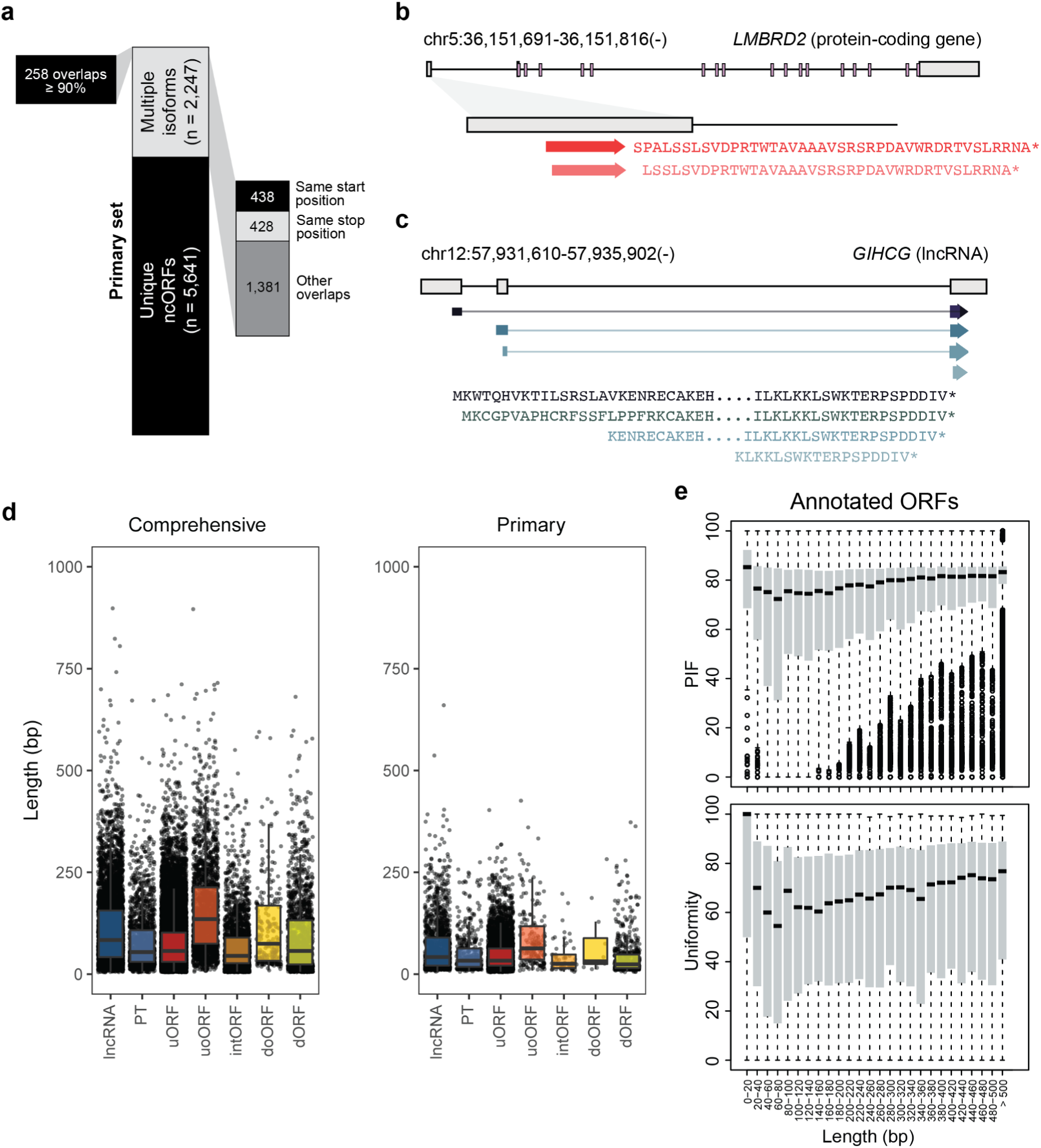
(a) Proportion and number of ncORFs in the Primary set, categorized based on the presence or absence of additional ncORF isoforms. For the set of 2,247 ncORFs associated with multiple isoforms, the number of overlapping isoforms (≥90% codon overlap with the shortest isoform) is indicated, as this threshold was used in the first catalog to collapse ncORFs. Additionally, all 2,247 ncORFs with multiple isoforms are further stratified by the type of overlap (same start position, same stop position, other overlaps). **(b)** Two overlapping uORF variants located in the *LMBRD2* gene have near-cognate initiation codons. **(c)** A translated region located in the *GIHCG* lncRNA has four different predicted initiation codons utilizing either an AUG initiation codon or a near-cognate initiation codon. **(d)** Boxplot showing length distribution of ncORFs across different ncORF types in the Comprehensive set and Primary set. **(e)** Boxplot showing PIF and Uniformity score distribution across length bins of annotated ORFs.

